# A chromosome-level genome assembly of *Drosophila madeirensis*, a fruit fly species endemic to the island of Madeira

**DOI:** 10.1101/2024.02.21.581469

**Authors:** Kenta Tomihara, Ana Llopart, Daisuke Yamamoto

**Author notes:** Corresponding authors (KT); (DY).

## Abstract

*Drosophila subobscura* is distributed across Europe, the Near East, and the Americas, while its sister species, *D. madeirensis*, is endemic to the island of Madeira in the Atlantic Ocean. *D. subobscura* is known for its strict light-dependence in mating and its unique courtship displays, including nuptial gift giving. *D. subobscura* has also attracted the interest of researchers because of its abundant variations in chromosomal polymorphisms correlated to the latitude and season, which have been used as a tool to track global climate warming. Although *D. madeirensis* can be an important resource for understanding the evolutionary underpinning of these genetic characteristics of *D. subobscura*, little work has been done on the biology of this species. Here, we used a HiFi long-read sequencing dataset to produce a *de novo* genome assembly for *D. madeirensis*. This assembly comprises a total of 111 contigs spanning 135.5 Mb, and has an N50 of 24.2 Mb and a BUSCO completeness score of 98.6%. Each of the six chromosomes of *D. madeirensis* consisted of a single contig. Breakpoints of the chromosomal inversions between *D. subobscura* and *D. madeirensis* were characterized using this genome assembly, updating some of the previously identified locations.

## Introduction

The genus *Drosophila* contains over 1,600 species (O’Grady and DeSalle 2018), which are highly divergent in morphology and behavior (Kopp *et al*. 2000; Prud’homme *et al*. 2006; Tanaka *et al*. 2009; Baker *et al*. 2023). Among these, *D. melanogaster* is one of the best-studied organisms in the animal kingdom: a large collection of mutants and genetically modified fly stocks as well as sophisticated genetic techniques have made it an unparalleled model for studies in all biological disciplines. Because it can be genetically modified in numerous ways, the functions of a large number of genes have been unveiled to date. Thus, comparative approaches with the members of the genus *Drosophila* will benefit enormously from the knowledge accumulated by the studies in *D. melanogaster. D. subobscura* offers a good starting point for such comparative approaches because classic genetics at the chromosomal level as well as modern molecular genetics including genome sequence data are publicly available as the potential substrate for addressing specific scientific questions.

*D. subobscura* was originally found around Europe (Buzzati-Traverso and Scossiroli 1955) and was more recently introduced to the Americas (Prevosti *et al*. 1988; Ayala *et al*. 1989). *D. subobscura* is known for a few unique displays in courtship behavior, such as nuptial gift giving (Steele 1986b, 1986a; Tanaka *et al*. 2017), an absolute light requirement for mating (Philip *et al*. 1944; Wallace and Dobzhansky 1946; Keesey *et al*. 2020), an absence of courtship songs (Ewing and Bennet-Clark 1968), and monoandry (Maynard Smith 1956; Fisher *et al*. 2013). The karyotype of *D. subobscura* consists of five large chromosomes named O, U, J, A, and E, and a small dot chromosome, and many variations in the sequence arrangement have been found across all five large chromosomes (reviewed in Sperlich and Pfriem 1986). Inversion polymorphisms show adaptive variation patterns across latitudes (Ayala *et al*. 2011) and seasons (Rodríguez-Trelles *et al*. 1996; Rodríguez-Trelles 2003). Thus, chromosomal inversion polymorphisms of *D. subobscura* have been used to track global climate warming (Rodríguez-Trelles and Rodríguez 1998; Balanyá *et al*. 2006). Recent long read-based whole genome sequencing helped unravel the link between inversions and seasonal adaptation in *D. subobscura* (Karageorgiou *et al*. 2020).

The *subobscura* subgroup consists of three species, namely, *D. subobscura, D. madeirensis* and *D. guanche* (Figure 1A) (Barrio *et al*. 1994; Barrio and Ayala 1997; Gao *et al*. 2007). Among them, *D. subobscura* and *D. madeirensis* are particularly closely related and are believed to have diverged allopatrically about 0.6–1.0 million years ago (Ramos-Onsins *et al*. 1998; Herrig *et al*. 2014). *D. madeirensis* is endemic to Madeira, an island in the Atlantic Ocean located approximately 580 km west of Morocco (Figure 1B) (Monclús 1984). Under laboratory conditions, *D. subobscura* and *D. madeirensis* can mate and some of their F_1_ hybrids are fertile (summarized in Table S1). In nature, limited gene flow was detected between *D. subobscura* on Madeira and *D. madeirensis* (Herrig *et al*. 2014). The existence of such a close yet separate outgroup will greatly facilitate understanding of the evolutionary basis for the emergence of distinctly different phenotypic characteristics among the sister species. However, while a substantial body of knowledge has been accumulated on the biology of *D. subobscura*, work on *D. madeirensis* has been sparse. In the present study, a highly complete and contiguous *de novo* genome assembly for *D. madeirensis* was constructed using HiFi long-read sequencing, which will serve as an important resource for future studies in all biology disciplines in this species subgroup.

**Figure 1.**
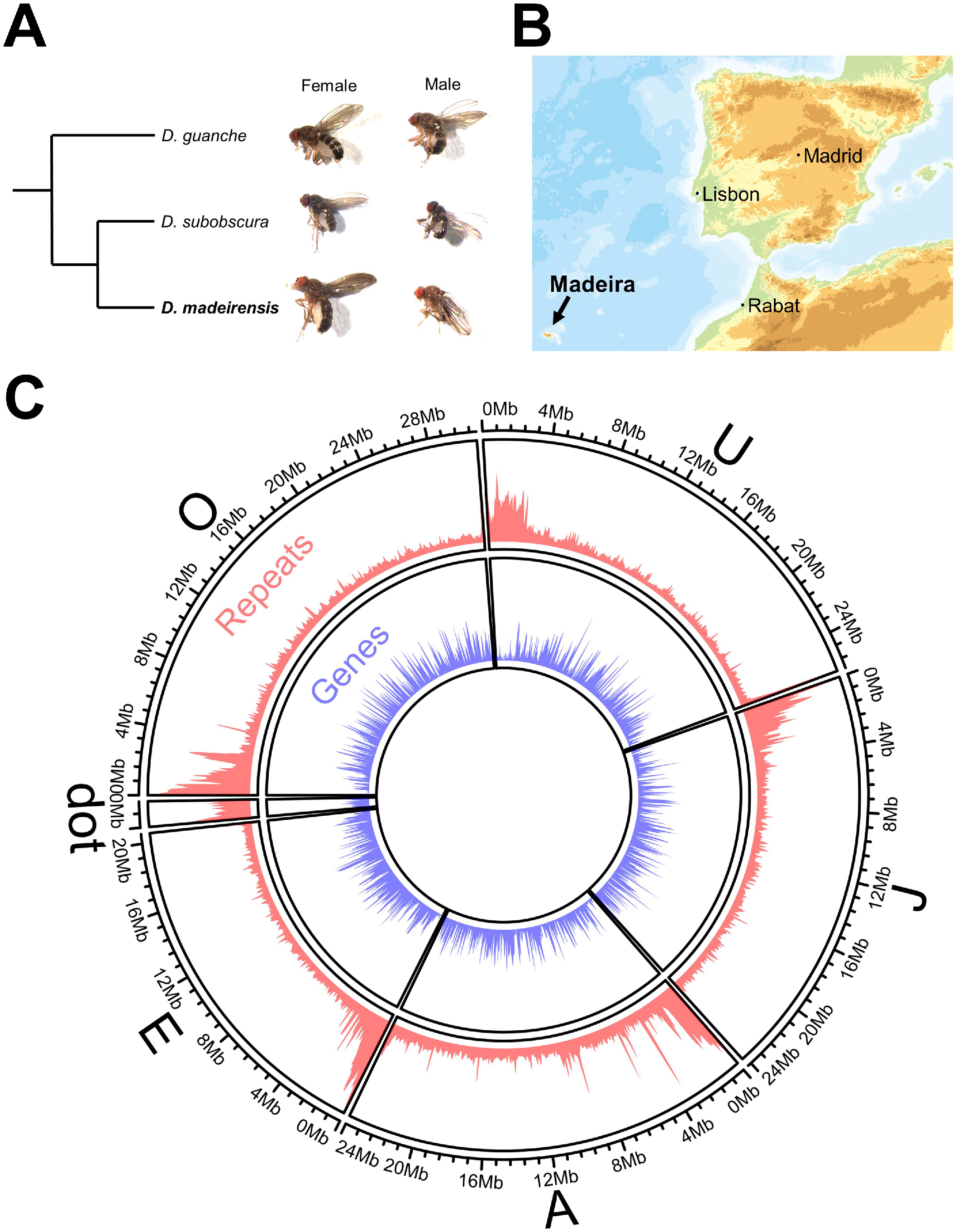
Overview of *D. madeirensis* and its genome. (A) A phylogenetic tree and pictures of fly samples of the species belonging to the *D. subobscura* subgroup. (B) Distribution of *D. madeirensis. D. madeirensis* is endemic to the island of Madeira in the Atlantic Ocean. The map was modified from the digital map published by the Geospatial Information Authority of Japan. (C) Representation of the distribution of genes and repeat sequences on each *D. madeirensis* assembled chromosome. Chromosomes are ordered by their size. The proportion of elements per each 100-kb nonoverlapping window is plotted as a histogram. The y-axis range is set to 0–1. The circular genome map was produced using the circlize R package (Gu *et al*. 2014).

## Materials and Methods

### Insects

The *D. madeirensis* strain RF1 was reared at 18ºC on standard cornmeal-yeast-agar medium, and kept in an incubator on a 12-h light/dark cycle. This strain was established from a single mated female (i.e., isofemale line) collected in Ribeiro Frio, Madeira (Herrig *et al*. 2014).

### Genome extraction and sequencing

Ten adult females were collected in one tube, and the genomic DNA was extracted using Genomic-tip 20/G (QIAGEN, Hilden, Germany) according to the manufacturer’s protocol. A library for HiFi sequencing was prepared using SMRTbell Prep Kit 3.0 (Pacific Biosciences, San Diego, CA), and sequencing was performed with Pacbio Sequel II (Pacific Biosciences). Library preparation and HiFi genome sequencing were performed by Takara Bio (Shiga, Japan). A library for short-read sequencing was prepared using an NEBNext Ultra II DNA Library Prep kit (New England Biolabs, Beverly, MA) and sequenced using a NovaSeq 6000 (Illumina, San Diego, CA) with 150 bp paired-end reads. Library preparation and genome sequencing for short-read sequencing were performed by Novogene (Beijing, China).

### Genome size estimation

Low-quality reads and adapter sequences were removed from raw short reads using Trim Galore! (https://www.bioinformatics.babraham.ac.uk/projects/trim_galore/) (Quality Phred score cutoff: 20, Maximum trimming error rate: 0.1, Minimum required adapter overlap: 1 bp, Minimum required sequence length for both reads before a sequence pair gets removed: 20 bp). The optimal k-mer length for this dataset was estimated by KmerGenie (Chikhi and Medvedev 2014), and then the genome size of *D. madeirensis* was estimated using this k-mer value (k-mer = 111).

### Genome assembly

HiFi reads from PacBio Sequel II were assembled with Hifiasm (Cheng *et al*. 2021), HiCanu (Nurk *et al*. 2020), PacBio’s Improved Phased (IPA; https://github.com/PacificBiosciences/pbipa), Flye (Kolmogorov *et al*. 2019), and NextDenovo (Hu *et al*. 2023). The options used in the assembler tools are listed in Table S2. Assembly statistics and completeness were calculated by QUAST-LG (Mikheenko *et al*. 2018) and BUSCO (Waterhouse *et al*. 2018), respectively. In BUSCO, consensus sequences were searched against the diptera_odb10 dataset. Because NextDenovo assembled most of the nuclear genome and the genome size matched that estimated by the k-mer method, we used this assembly for further analyses (see Results and Discussion).

The repertory of repetitive elements in the genome sequence assembled by NextDenovo was obtained using RepeatModeler (Flynn *et al*. 2020). Based on this repertory, repetitive sequences were masked from the assembled genome by RepeatMasker (http://www.repeatmasker.org). For each masked contig, BLASTn was performed against the NCBI nt database and the sequence with the highest homology was obtained (options: -task megablast -evalue 1e-10 -soft_masking true). Contigs that were assigned to the sequences derived from bacterial species were removed from the genome assembly. To calculate coverage, HiFi reads were mapped to this assembly using minimap2 (Li 2018) with the map-hifi option.

### RNA extraction, sequencing, and gene annotation

Total RNA was extracted from embryos, larvae, pupae, and adults (Table S3) using TRIzol reagent (Thermo Fisher Scientific, Waltham, MA) according to the manufacturer’s protocol. Poly-A selected libraries were prepared with a NEBNext Ultra II Directional RNA Library Prep Kit (New England Biolabs) and sequenced using the NovaSeq 6000 (Ilumina) with 150 bp paired-end reads. Library preparation and sequencing were performed by Novogene.

Low-quality reads and adapter sequences were removed from raw reads using Trim Galore! with the same options as for genomic short reads. Gene models were predicted by BRAKER3 (Gabriel *et al*. 2023) using a repeat-masked genome assembly, trimmed RNA-seq reads, and a database of protein families across arthropods obtained from OrthoDB (Kuznetsov *et al*. 2023). The gene models were annotated using a DIAMOND BLASTp search (Buchfink *et al*. 2021) against NCBI RefSeq database.

### Alignment of *D. madeirensis* and *D. subobscura* genome sequences

The *D. madeirensis* genome was aligned to the *D. subobscura* reference genome (Bracewell *et al*. 2019) by minimap2 (Li 2018) with the asm5 option. Harr plots were drawn by pafr (https://github.com/dwinter/pafr) and a custom R script.

## Results and Discussion

### Genome assembly

With the HiFi dataset, we compared the performance of five assemblers: Hifiasm (Cheng *et al*. 2021), HiCanu (Nurk *et al*. 2020), IPA (https://github.com/PacificBiosciences/pbipa), Flye (Kolmogorov *et al*. 2019), and NextDenovo (Hu *et al*. 2023) (Table 1). There were large differences in contig continuity. The largest contigs in Hifiasm and HiCanu were assembled to 43.2 Mb, whereas those of NextDenovo, IPA, and Flye were assembled to 30.8, 30.6 and 19.2 Mb, respectively. NextDenovo and HiCanu showed the two highest N50 values (24.2 and 21.2 Mb, respectively) among the assemblers (3.3–16.1 Mb). Overall, Hifiasm, HiCanu, and NextDenovo performed better than the rest of the assemblers in terms of contig continuity. The total contig lengths also varied substantially according to the assembler (Table 1; 146.3–328.2 Mb). Using the k-mer method, we evaluated the genome size of *D. madeirensis* to be 140.6 Mb. A previous work estimated the genome size of *D. subobscura* to be about 137 Mb by k-mer and 148 Mb by flow cytometry (Karageorgiou *et al*. 2019), in line with our estimation for the *D. madeirensis* genome. Taking these results together, we consider that the size of the total contig lengths was correctly estimated by NextDenovo but overestimated by the other four assemblers. Therefore, we conclude that the genome sequence assembled by NextDenovo would be the most reliable in regard to both the contig continuity and genome size.

**Table 1.** Statistics of genomes assembled by five different assemblers.

| Assembler | QUAST-LG |  |  |  |  | BUSCO (3285 groups searched) |  |  |  |
| --- | --- | --- | --- | --- | --- | --- | --- | --- | --- |
|  | Number of contigs | Total length (bp) | Largest contig (bp) | N50 (bp) | L50 | Complete and single-copy | Complete and duplicated | Fragmented | Missing |
| Hifiasm | 226 | 292,387,389 | 43,242,118 | 21,175,663 | 5 | 3,203 | 54 | 17 | 11 |
| HiCanu | 1,478 | 328,219,170 | 43,238,710 | 16,072,932 | 8 | 3,081 | 176 | 17 | 11 |
| IPA | 110 | 174,598,151 | 30,560,687 | 9,811,834 | 6 | 3,240 | 17 | 17 | 11 |
| Flye | 747 | 273,503,793 | 19,173,260 | 3,263,947 | 16 | 3,172 | 85 | 17 | 11 |
| NextDenovo | 146 | 146,286,141 | 30,799,563 | 24,201,710 | 3 | 3,241 | 18 | 15 | 11 |
| Final assembly (contaminants removed) | 111 | 135,508,623 | 30,799,563 | 24,201,710 | 3 | 3,240 | 18 | 16 | 11 |

To identify possible contaminants in the contigs obtained with NextDenovo, we performed BLASTn searches against the NCBI nt genome database. Out of 146 contigs, 24 were assigned to *Gluconobacter* (5.5 Mb in total), 9 were assigned to *Serratia* (5.1 Mb), 1 was assigned to *Klebsiella* (85 kb), and 1 was assigned to *Komagataeibacter* (45 kb) bacterial species. All these bacteria are inhabitants of the *Drosophila* gut (Staubach *et al*. 2013; Chaston *et al*. 2016; Ramírez-Camejo *et al*. 2017; Gao *et al*. 2023). We analyzed the sequence after removing these contigs derived from bacteria. The average coverage of HiFi reads across this assembly is 89.67×. This assembly comprised a total of 111 contigs spanning 135.5 Mb and had an N50 of 24.2 Mb (Table 1). Of the 3,285 dipteran BUSCOs, 3,240 (98.6%) were found to be complete and single-copy (Table 1). All six chromosomes of *D. madeirensis* were assembled into a single contig (Figure 1).

### Gene annotation

Repeat sequences were predicted using RepeatMasker, which covered 19.5% of the whole genome (Figure 1C). Using RNA-seq data and a protein dataset across arthropods obtained from OrthoDB (Kuznetsov et al. 2023), BRAKER3 (Gabriel et al. 2023) annotated 13,789 protein-coding genes in the *D. madeirensis* genome, which would potentially produce 17,455 unique proteins (Figure 1C).

Centromeres in many multicellular eukaryotes consist of highly repetitive DNA sequences (Melters *et al*. 2013). In *D. subobscura*, all five large chromosomes are telocentric (i.e., the centromeres are at the end of chromosomes) (Buzzati-Traverso and Scossiroli 1955; Bracewell *et al*. 2019). We found that repetitive DNA sequences in the *D. madeirensis* genome are concentrated at the end of chromosomes, while gene density was lower at the end of chromosomes (Figure 1C). This pattern is similar to that of *D. subobscura* (Bracewell *et al*. 2019), suggesting that *D. madeirensis* also has telocentric chromosomes.

### Chromosomal structures of *D. madeirensis* and *D. subobscura*

Figure 2 shows the genome alignment of *D. madeirensis* and the *D. subobscura* reference strain (14011-0131.10; Bracewell et al. 2019). All six *D. subobscura* chromosomes were uniquely aligned with the *D. madeirensis* chromosomes, demonstrating the contiguity and completeness of both assemblies. In previous studies, the chromosomal structure of *D. madeirensis* was referred to as O_ms_, U_1+2_, J_ST_, A_h1+h2+h3+5,_ and E_ST_, while that of the *D. subobscura* 14011-0131.10 strain was called O_ms+4_, U_1+2_, J_ST_, A_ST_, and E_ST_ (Karageorgiou *et al*. 2019, 2020). There are no obvious structural rearrangements of chromosomes U, J, and E between the two species, so that the same names are used to describe them. In contrast, there are large inversions on chromosomes A and O and a small additional inversion at the end of chromosome A of *D. madeirensis*, consistent with previous polytene chromosome analyses of the hybrids between *D. madeirensis* and *D. subobscura* (Krimbas and Loukas 1984; Papaceit and Prevosti 1989).

**Figure 2.**
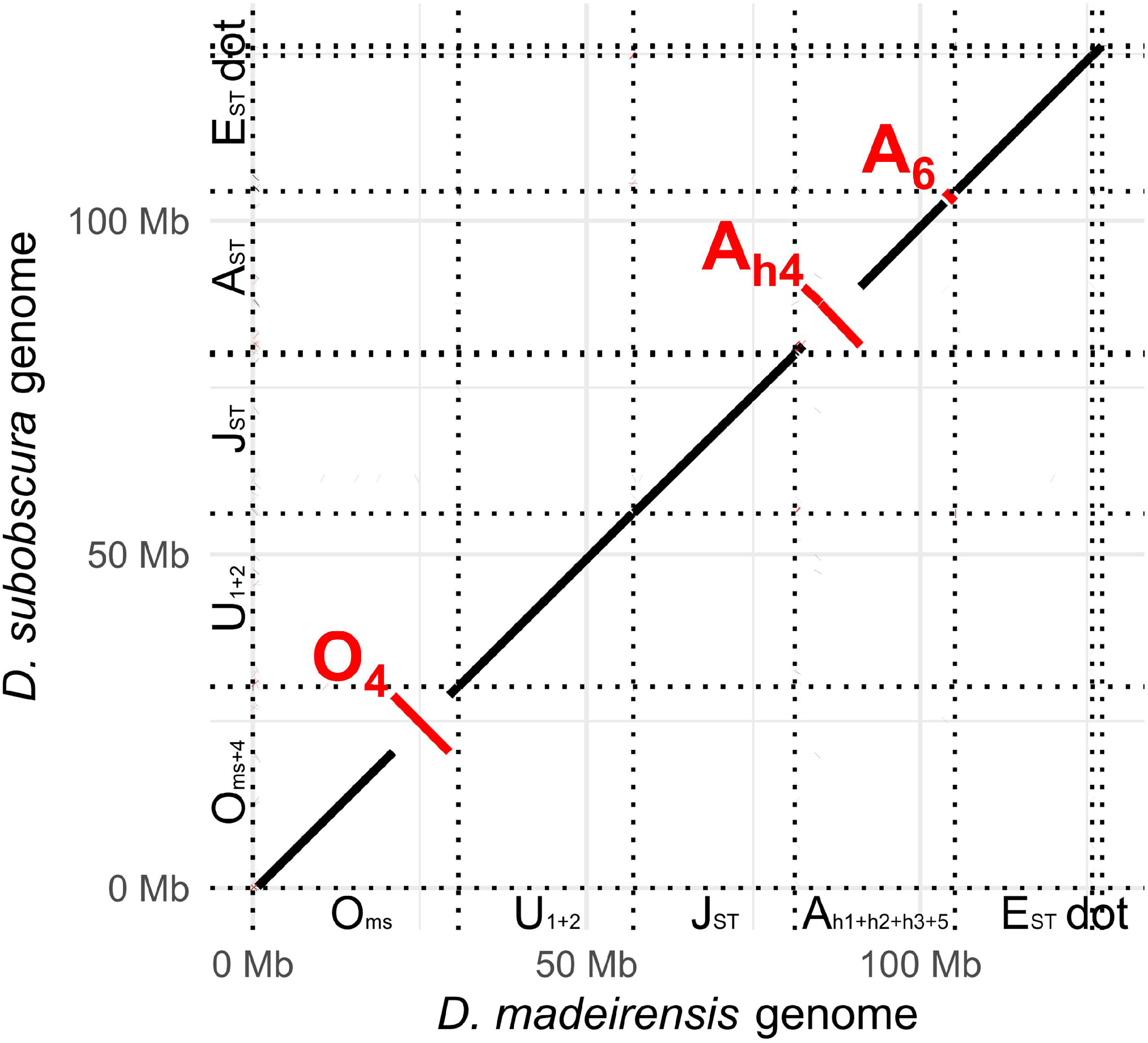
Harr-plot made with minimap2 (Li 2018) whole-genome alignment between *D. madeirensis* and *D. subobscura* genome sequences. Sequences aligned in forward and reverse orientations are represented by black and red lines, respectively. Inversions O_4_, A_h4_, and A_6_ are indicated by red letters.

Breakpoints of inversion O_4_ found in the *D. subobscura* O_ms+4_ chromosome were previously characterized using different *D. subobscura* strains: the proximal O_4_ breakpoint contains two truncated genes, i.e., *Peroxidase* (*Pxd*) and *CG5225*, while the distal O_4_ inversion breakpoint contains another two, also truncated, genes, i.e., *SET domain containing 8* (*Set8*) and *ATP-dependent chromatin assembly factor large subunit* (*Acf*) (Puerma *et al*. 2016). Here the gene names are based on the *Drosophila melanogaster* ortholog. Our alignment of *D. subobscura* and *D. madeirensis* genomes confirmed that the O_4_ inversion occurred at these breakpoints (Figure 3A, Table S4).

**Figure 3.**
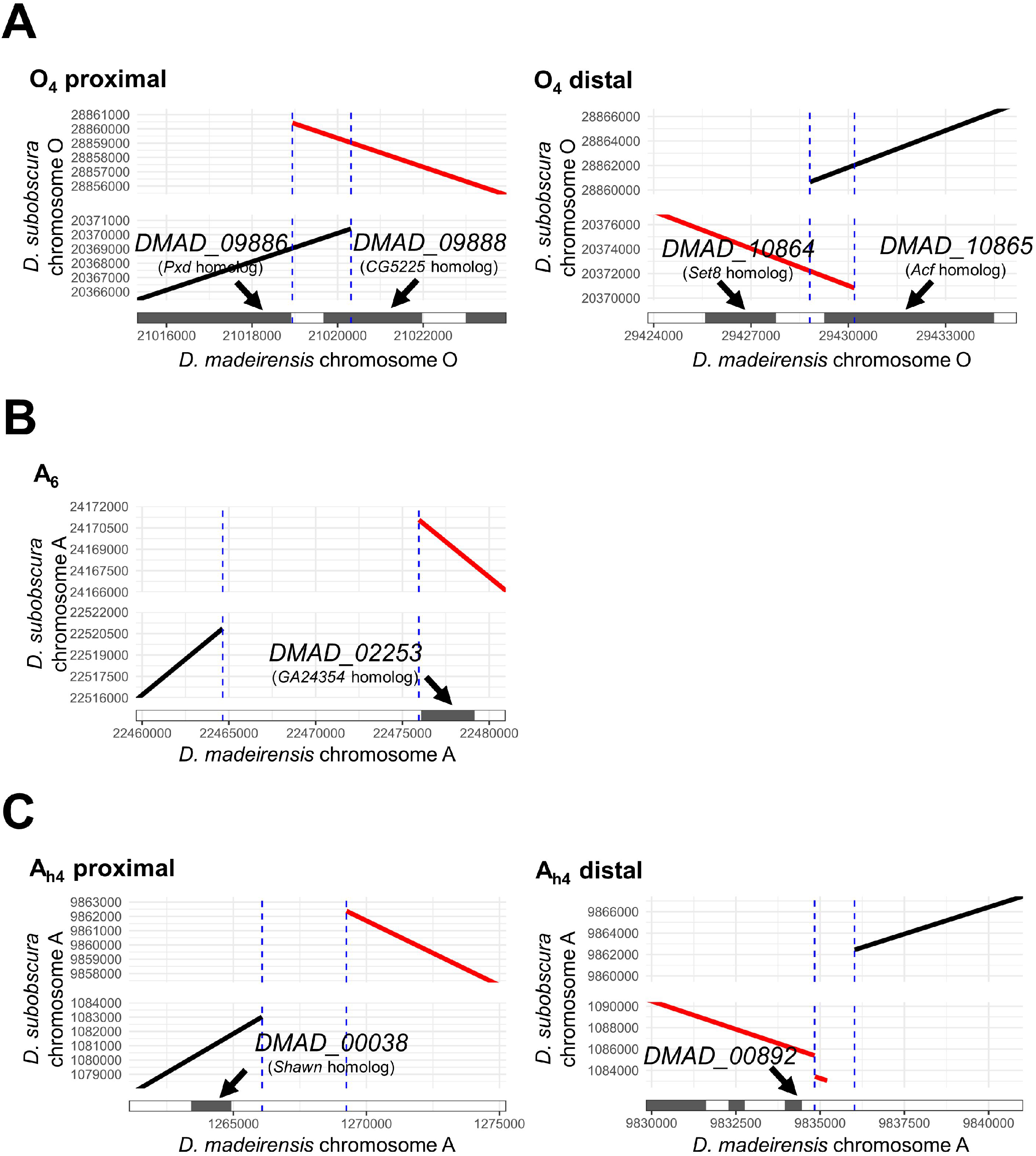
Detailed characterization of inversion breakpoints. Harr plots of *D. madeirensis* and *D. subobscura* genome sequences at the O_4_ (A), A_6_ (B), and A_h4_ (C) breakpoints are shown. Sequences aligned in forward and reverse orientations are represented by black and red lines, respectively. The blue dotted lines indicate breakpoints of the inversions. Each filled rectangle represents a *D. madeirensis* gene model.

A_h1+h2+h3+5_ and A_ST_ were predicted to have two inversions, namely A_h4_ and A_6_ (Karageorgiou *et al*. 2019). Because A_6_ is a terminal inversion, it has only one breakpoint. By designing molecular markers on *D. subobscura* and *D. guanche* genomes, Orengo et al. (2019) mapped the A_6_ breakpoint to a ~20-kb long region flanked by genes homologous to *D. pseudoobscura GA22805* and *GA24354* (note that A_6_ and A_h4_ were referred to as A_f6_ and A_f1_ in their study, respectively). Our alignment of the *D. subobscura* and *D. madeirensis* genomes further narrowed the A_6_ breakpoint to a 11.3 kb region near *DMAD_02253*, a gene homologous to *GA24354* (Figure 3B, Table S4), which supports the result of the previous study. Orengo et al. (2019) also determined the proximal and distal breakpoints of A_h4_: a ~44-kb region flanked by the genes homologous to *D. pseudoobscura GA14783* (corresponding to the *D. madeirensis* gene *DMAD_00872*) and *GA13678* (*DMAD_00079*), and a ~33-kb region flanked by *GA17070* (*DMAD_00099*) and *GA15499* (*DMAD_00868*), respectively. However, our alignment of the *D. subobscura* and *D. madeirensis* genomes gave a different result. We located the proximal A_h4_ breakpoint on a 3.2 kb region near *DMAD_00038*, and the distal break point on a 1.2 kb region near *DMAD_00892* (Figure 3C, Table S4). The proximal and distal A_h4_ breakpoints determined in this study were about 521 and 87 kb away from the closest genes characterized by Orengo et al. (2019), respectively (Figure S1, Table S4). We found many inversions on the A chromosome that had occurred after the separation of *D. guanche* from the clade of *D. subobscura*/*D. madeirensis* (Figure S2), and this may have complicated the previous mapping of the A_h4_ breakpoints, which relied on molecular markers designed solely based on the genomic information of *D. guanche* and *D. subobscura* (Orengo *et al*. 2019). These considerations highlight the importance of using closely related lineages with fewer chromosomal rearrangements in mapping chromosomal breakpoints.

The comprehensive knowledge of the *D. madeirensis* genome obtained in this study will provide a solid basis for future comparative analyses of the diverged phenotypic traits found among the three members of the *D. subobscura* species subgroup.

## Data Availability Statement

The raw sequence data have been submitted to DDBJ under accession numbers DRR528066 (genomic Hifi reads), DRR528092 (genomic short reads), and DRR528093–528099 (RNA-seq reads). The *D. madeirensis* genome assembly and gene models have been submitted to DDBJ under accession number PRJDB17459.

## Acknowledgments

We thank Ryoya Tanaka for useful comments. We also thank Manami Adachi for secretarial assistance.

## Conflict of Interest

The authors declare that they have no conflict of interest.

## Funder Information

This work was supported by a JSPS Grant-in-Aid (no. JP23KJ2210) to KT and a MEXT Grant-in-Aid (no. JP21H04790) to DY.

## Author Contributions

KT and DY designed the study. KT conducted most of the experiments and analyses. AL provided the insects used in this study. KT wrote the manuscript. KT and DY reviewed and edited the manuscript. All authors approved the final version of the manuscript and agree to be accountable for all aspects of the work.

